# Personalized genetic assessment of age associated Alzheimer’s disease risk

**DOI:** 10.1101/074864

**Authors:** Rahul S. Desikan, Chun Chieh Fan, Yunpeng Wang, Andrew J. Schork, Howard J. Cabral, L. Adrienne Cupples, Wesley K. Thompson, Lilah Besser, Walter A. Kukull, Dominic Holland, Chi-Hua Chen, James B. Brewer, David S. Karow, Karolina Kauppi, Aree Witoelar, Celeste M. Karch, Luke W. Bonham, Jennifer S. Yokoyama, Howard J. Rosen, Bruce L. Miller, William P. Dillon, David M. Wilson, Christopher P. Hess, Margaret Pericak-Vance, Jonathan L. Haines, Lindsay A. Farrer, Richard Mayeux, John Hardy, Alison M. Goate, Bradley T. Hyman, Gerard D. Schellenberg, Linda K. McEvoy, Ole A. Andreassen, Anders M. Dale, for the ADNI and ADGC investigators

## Abstract

**Importance:** Identifying individuals at risk for developing Alzheimer’s disease (AD) is of utmost importance. Although genetic studies have identified *APOE* and other AD associated single nucleotide polymorphisms (SNPs), genetic information has not been integrated into an epidemiological framework for personalized risk prediction.

**Objective:** To develop, replicate and validate a novel polygenic hazard score for predicting age-specific risk for AD.

**Setting:** Multi-center, multi-cohort genetic and clinical data.

**Participants:** We assessed genetic data from 17,008 AD patients and 37,154 controls from the International Genetics of Alzheimer’s Project (IGAP), and 6,409 AD patients and 9,386 older controls from Phase 1 Alzheimer’s Disease Genetics Consortium (ADGC). As independent replication and validation cohorts, we also evaluated genetic, neuroimaging, neuropathologic, CSF and clinical data from ADGC Phase 2, National Institute of Aging Alzheimer’s Disease Center (NIA ADC) and Alzheimer’s Disease Neuroimaging Initiative (ADNI) (total n = 20,680)

**Main Outcome(s) and Measure(s):** Use the IGAP cohort to first identify AD associated SNPs (at p < 10^-5^). Next, integrate these AD associated SNPs into a Cox proportional hazards model using ADGC phase 1 genetic data, providing a polygenic hazard score (PHS) for each participant. Combine population based incidence rates, and genotype-derived PHS for each individual to derive estimates of instantaneous risk for developing AD, based on genotype and age. Finally, assess replication and validation of PHS in independent cohorts.

**Results:** Individuals in the highest PHS quantiles developed AD at a considerably lower age and had the highest yearly AD incidence rate. Among *APOE* ε3/3 individuals, PHS modified expected age of AD onset by more than 10 years between the lowest and highest deciles. In independent cohorts, PHS strongly predicted empirical age of AD onset (p = 1.1 x 10^-26^), longitudinal progression from normal aging to AD (p = 1.54 x 10^-10^) and associated with markers of AD neurodegeneration.

**Conclusions:** We developed, replicated and validated a clinically usable PHS for quantifying individual differences in age-specific risk of AD. Beyond *APOE*, polygenic architecture plays an important role in modifying AD risk. Precise quantification of AD genetic risk will be useful for early diagnosis and therapeutic strategies.

## INTRODUCTION

Late onset Alzheimer’s disease (AD), the most common form of dementia, places a large emotional and economic burden on patients and society. With increasing health care expenditures among cognitively impaired elderly^1^, identifying individuals at risk for developing AD is of utmost importance for potential preventative and therapeutic strategies. Inheritance of the ε4 allele of apolipoprotein E (*APOE*) on chromosome 19q13 is the most significant risk factor for developing late-onset AD.^2^ *APOE* ε4 has a dose dependent effect on age of onset, increases AD risk three-fold in heterozygotes and fifteen-fold in homozygotes, and is implicated in 20-25% of patients with AD.^3^

In addition to *APOE*, recent genome-wide association studies (GWAS) have identified numerous AD associated single nucleotide polymorphisms (SNPs), most of which have a small effect on disease risk.^4-5^ Although no single polymorphism may be informative clinically, a combination of *APOE* and non-*APOE* SNPs may help identify older individuals at increased risk for AD. Despite the detection of novel AD associated genes, GWAS findings have not yet been incorporated into a genetic epidemiology framework for individualized risk prediction.

Building on a prior approach evaluating GWAS-detected genetic variants for disease prediction^7^ and using a survival analysis framework, we tested the feasibility of combining AD associated SNPs and *APOE* status into a continuous measure ‘polygenic hazard score’ (PHS) for predicting the age-specific risk for developing AD. We assessed replication and validation of the PHS using several independent cohorts.

## METHODS

### Participant Samples

#### IGAP

To select AD associated SNPs, we evaluated publicly available AD GWAS summary statistic data (p-values and odds ratios) from the International Genomics of Alzheimer’s Disease Project (IGAP Stage 1, for additional details see Supplemental Information and reference 4). We used IGAP Stage 1 data, consisting of 17,008 AD cases and 37,154 controls, for selecting AD associated SNPs (for a description of the AD cases and controls within the IGAP Stage 1 sub-studies, please see Table 1 and reference 4).

#### ADGC

To develop the survival model for the polygenic hazard scores (PHS), we first evaluated age of onset and raw genotype data from 6,409 patients with clinically diagnosed AD and 9,386 cognitively normal older individuals provided by the Alzheimer’s Disease Genetics Consortium (ADGC, Phase 1, a subset of the IGAP dataset), excluding individuals from the National Institute of Aging Alzheimer’s Disease Center (NIA ADC) samples and Alzheimer’s Disease Neuroimaging Initiative (ADNI). To evaluate replication of PHS, we used an independent sample of 6,984 AD patients and 10,972 cognitively normal older individuals from the ADGC Phase 2 cohort (Table 1). A detailed description of the genotype and phenotype data within the ADGC datasets has been described in detail elsewhere.^7,24^ Briefly, the ADGC Phase 1 and 2 datasets consist of multi-center, case-control, prospective, and family-based sub-studies of Caucasian participants with AD occurrence after age 60. Participants with autosomal dominant (*APP*, *PSEN1* and *PSEN2*) mutations were excluded. All participants were genotyped using commercially available high-density SNP microarrays from Illumina or Affymetrix. Clinical diagnosis of AD within the ADGC sub-studies was established using NINCDS/ADRDA criteria for definite, probable or possible AD. ^8^ For most participants, age of AD onset was obtained from medical records and defined as the age when AD symptoms manifested, as reported by the participant or an informant. For participants lacking age of onset, age at ascertainment was used. Patients with an age-at-onset or age-at-death less than 60 years, and Caucasians of European ancestry were excluded from the analyses. For additional details regarding the ADGC datasets, please see references 7 and 24.

**Table 1.** Demographic data for AD patients and older controls.

|  | IGAP<br>AD<br>patients | IGAP<br>older<br>controls | ADGC<br>Phase 1<br>AD<br>patients | ADGC<br>Phase 1<br>older<br>controls | ADGC<br>Phase<br>2 AD<br>patients | ADGC<br>Phase<br>2 older<br>controls |
| --- | --- | --- | --- | --- | --- | --- |
| **Total N** | 17,008 | 37,154 | 6,409 | 9,386 | 6,984 | 10,972 |
| **Mean age**<br>**(SD) of**<br>**onset**<br>**(cases) or**<br>**assessment**<br>**(controls)** | 74.7<br>(8.0) | 68.6<br>(8.5) | 74.7<br>(7.7) | 76.4<br>(8.1) | 73.6<br>(7.3) | 75.7<br>(8.6) |
| **%**<br>**Female** | 63 | 57 | 61 | 59 | 57.6 | 60.7 |
| **% *APOE***<br>**ε4**<br>**carriers** | 59.0 | 25.4 | 51.6 | 26.7 | 56.0 | 28.4 |

#### NIA ADC

To assess longitudinal prediction, we evaluated an ADGC-independent sample of 2,724 cognitively normal elderly individuals with at least 2 years of longitudinal clinical follow-up derived from the NIA funded ADCs (data collection coordinated by the National Alzheimer’s Coordinating Center). ^9^ To assess the relationship between polygenic risk and neuropathology, we assessed 2,960 participants from the NIA ADC samples with genotype and neuropathological evaluations. For the neuropathological variables, we examined the Braak stage for neurofibrillary tangles (NFTs) (0: none; I-II: entorhinal; III-IV: limbic, and V-VI: isocortical) ^10^ and the Consortium to Establish a Registry for Alzheimer’s Disease (CERAD) score for neuritic plaques (none/sparse, moderate, or frequent). ^11^

#### ADNI

To assess the relationship between polygenic risk and *in vivo* biomarkers, we evaluated an ADGC-independent sample of 692 older controls, mild cognitive impairment and AD participants from the ADNI (see Supplemental Methods). On a subset of ADNI1 participants with available genotype data, we evaluated baseline CSF levels of Aβ_1-42_ and total tau, as well as longitudinal clinical dementia rating-sum of box (CDR-SB) scores. In ADNI1 participants with available genotype and quality-assured baseline and follow-up MRI scans, we also assessed longitudinal sub-regional change in medial temporal lobe volume (atrophy) on 2471 serial T_1_-weighted MRI scans (for additional details see Supplemental Methods).

### Statistical Analysis

We followed three steps to derive the polygenic hazard scores (PHS) for predicting AD age of onset: 1) we defined the set of associated SNPs, 2) we estimated hazard ratios for polygenic profiles, and 3) we calculated individualized absolute hazards (see Supplemental Information for detailed description of these steps).

Using the IGAP Stage 1 summary statistics, we first identified a list of SNPs associated with increased risk for AD using significance threshold of p < 10^−5^. Next, we evaluated all IGAP-detected, AD-associated SNPs within the ADGC Phase 1 case-control dataset. Using a stepwise procedure in survival analysis, we delineated the final list of SNPs for constructing the polygenic hazard score. ^12−13^ In the Cox proportional hazard models, we identified the top AD-associated SNPs within the ADGC Phase 1 cohort (excluding NIA ADC and ADNI samples), while controlling for the effects of gender, *APOE* variants, and top five genetic principal components (to control for the effects of population stratification). We utilized age of AD onset and age of last clinical visit to estimate ‘age appropriate’ hazards ^14^ and derived a PHS for each participant. In each step of the stepwise procedure, the algorithm selected one SNP from the pool that most improved model prediction (i.e. minimizing the Martingale residuals); additional SNP inclusion that did not further minimize the residuals resulted in halting of the selection process. To prevent over-fitting in the training step, we used 1000x bootstrapping for model averaging and estimating the hazard ratios for each selected SNPs. We assessed the proportional hazard assumption in the final model using graphical comparisons.

To assess replication, we first examined whether the ADGC Phase 1 derived predicted PHSs could stratify individuals into different risk strata within the ADGC Phase 2 cohort. We next evaluated the relationship between predicted age of AD onset and the empirical/actual age of AD onset using cases from ADGC Phase 2. We binned risk strata into percentile bins and calculated the mean of actual age in that percentile as the empirical age of AD onset.

Because case-control samples cannot provide the proper baseline hazard, ^16^ we used the previously reported annualized incidence rates by age, estimated from the general United States of America (US) population. ^17^ For each participant, by combining the overall population-derived incidence rates ^17^ and genotype-derived PHS, we calculated an individual’s instantaneous risk for developing AD, based on their genotype and age (for additional details see Supplemental Information). To independently validate the predicted instantaneous risk, we evaluated longitudinal follow-up data from 2,724 cognitively normal older individuals from the NIA ADC with at least 2 years of clinical follow-up. We assessed the number of cognitively normal individuals progressing to AD as a function of the predicted PHS risk strata and examined whether the predicted PHS-derived incidence rate reflects the empirical/actual progression rate using a Cochran-Armitage trend test.

To assess validity, we examined the association between our PHS and established *in vivo* and pathologic markers of AD neurodegeneration. Using linear models, we assessed whether the PHS correlated with Braak stage for NFTs and CERAD score for neuritic plaques as well as CSF Aβ_1-42_, and CSF total tau. Using linear mixed effects models, we also investigated whether the PHS was associated with longitudinal CDR-SB score and volume loss within the entorhinal cortex and hippocampus. In all analyses, we co-varied for the effects of age and sex.

## RESULTS

### PHS: model development, relationship to APOE and independent replication

From the IGAP cohort, we found 1854 SNPs associated with increased risk for AD at a p < 10^-5^. Of these, using the Cox stepwise regression framework, we identified 31 SNPs, in addition to two *APOE* variants, within the ADGC cohort for inclusion into the polygenic model (Table 2). Figure 1 illustrates the relative risk for developing AD using the ADGC case/control Phase 1 cohort. The graphical comparisons among Kaplan-Meier estimations and Cox proportional hazard models indicate the proportional hazard assumption holds for the final model (Figure 1).

**Table 2.**
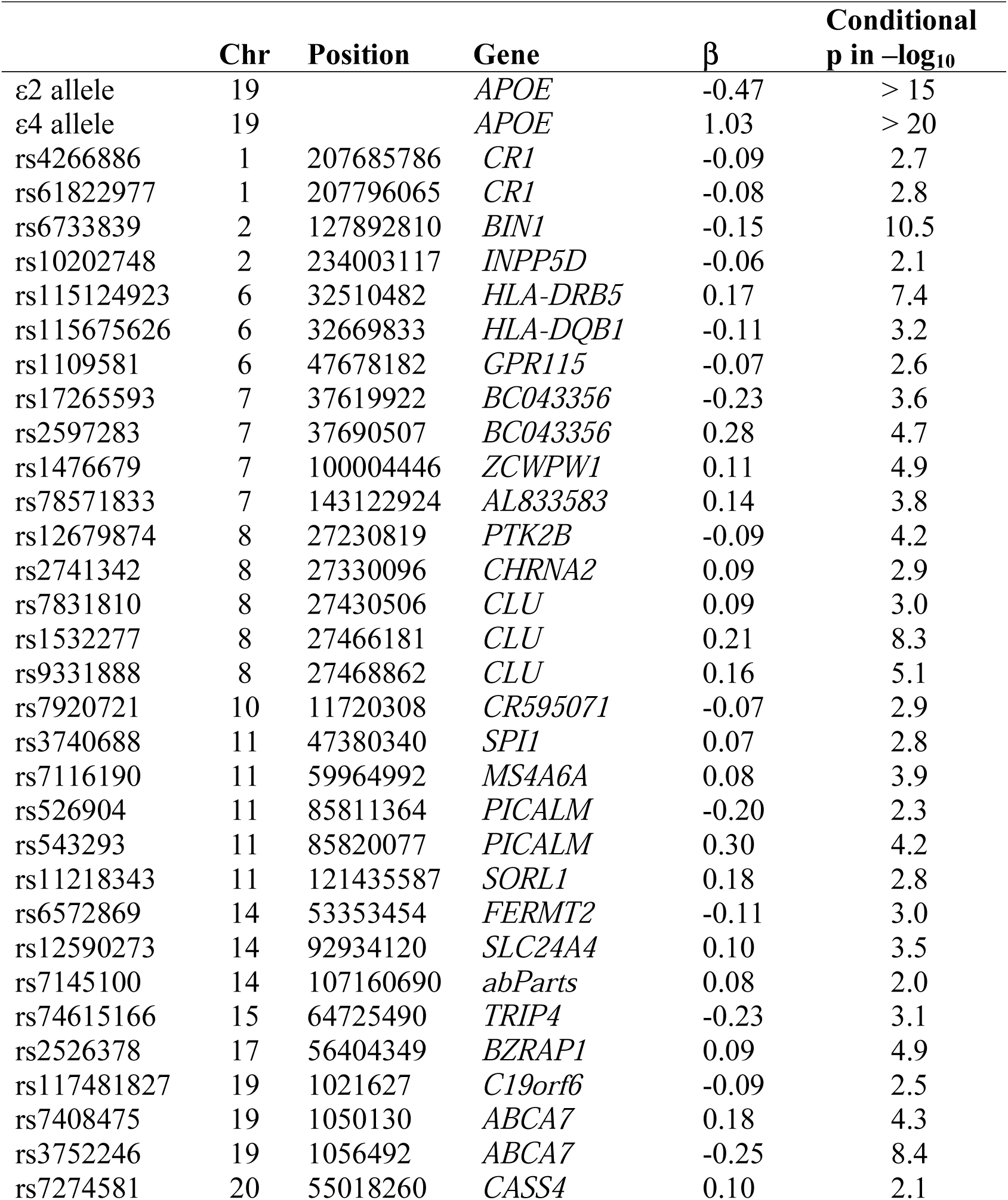
Selected 31 SNPs, their closest genes, hazard ratio estimations, and their conditional p values in the final joint model, after controlling for effects of gender and APOE variants.

**Figure 1.**
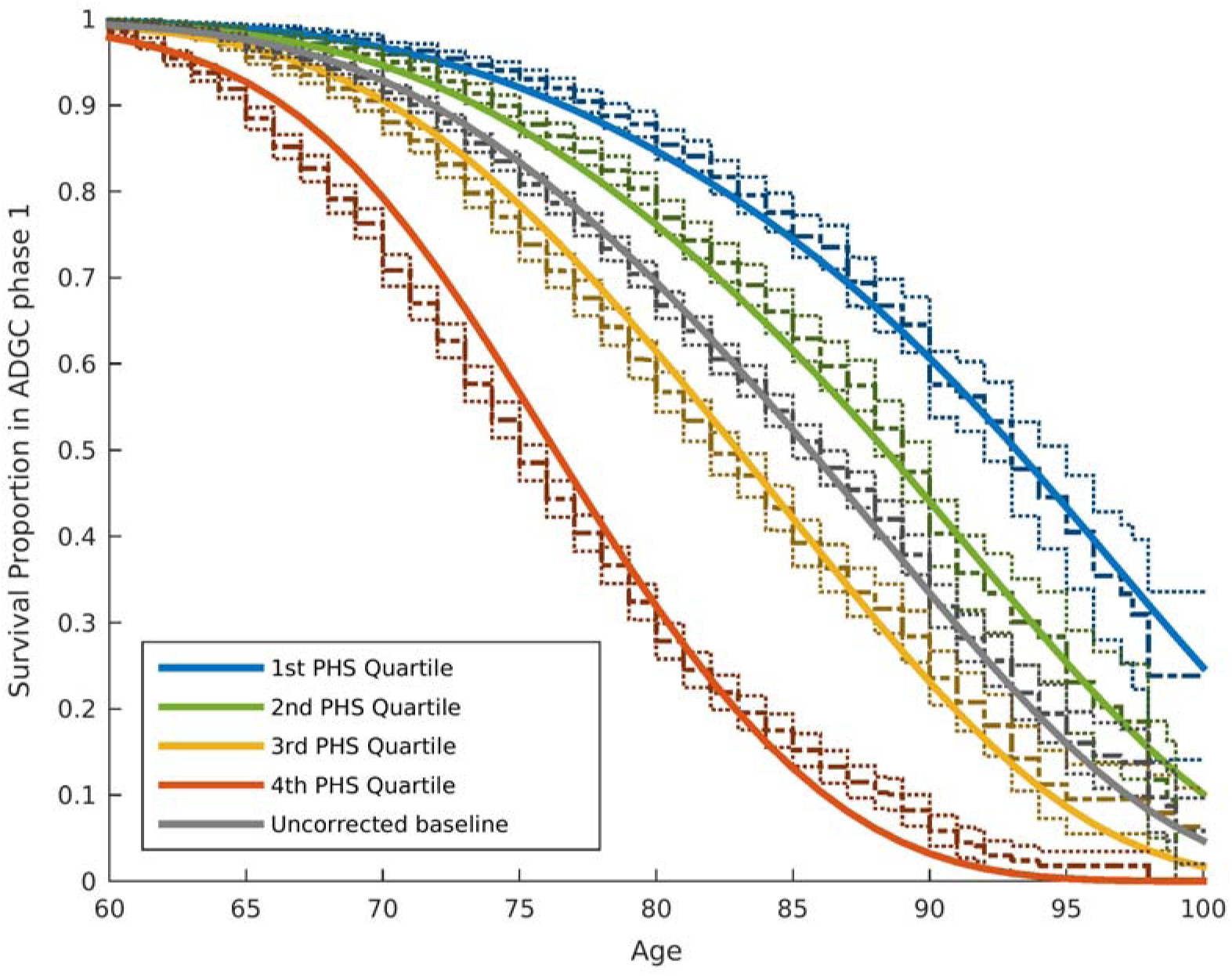
Kaplan-Meier estimates and Cox proportional model fits from the case-control ADGC phase 1 dataset, excluding NACC and ADNI samples. The proportional hazard assumptions were checked based on the graphical comparisons between Kaplan-Meier estimation and Cox proportional hazard models. 95% confidence intervals of Kaplan-Meier estimation are also demonstrated. The baseline hazard (gray line) in this model is based on the mean of ADGC data.

To quantify the additional prediction provided by polygenic information beyond *APOE*, we evaluated how PHS modulates age of AD onset in *APOE* ε3/3 individuals. Among these individuals, we found that age of AD onset can vary by more than 10 years, depending on polygenic risk. For example, for an *APOE* ε3/3 individual in the 10^th^ decile (top 10%) of PHS, at a survival proportion of 50%, the expected age for developing AD is approximately 84 years (Figure 2); however, for an *APOE* ε3/3 individual in the 1^st^ decile (bottom 10%) of PHS, the expected age of developing AD is approximately 95 years (Figure 2). Similarly, we also evaluated the relationship between PHS and the different *APOE* alleles (ε 2/3/4) (Supplemental Figure 1). These findings show that beyond *APOE*, the polygenic architecture plays an integral role in affecting AD risk.

**Figure 2.**
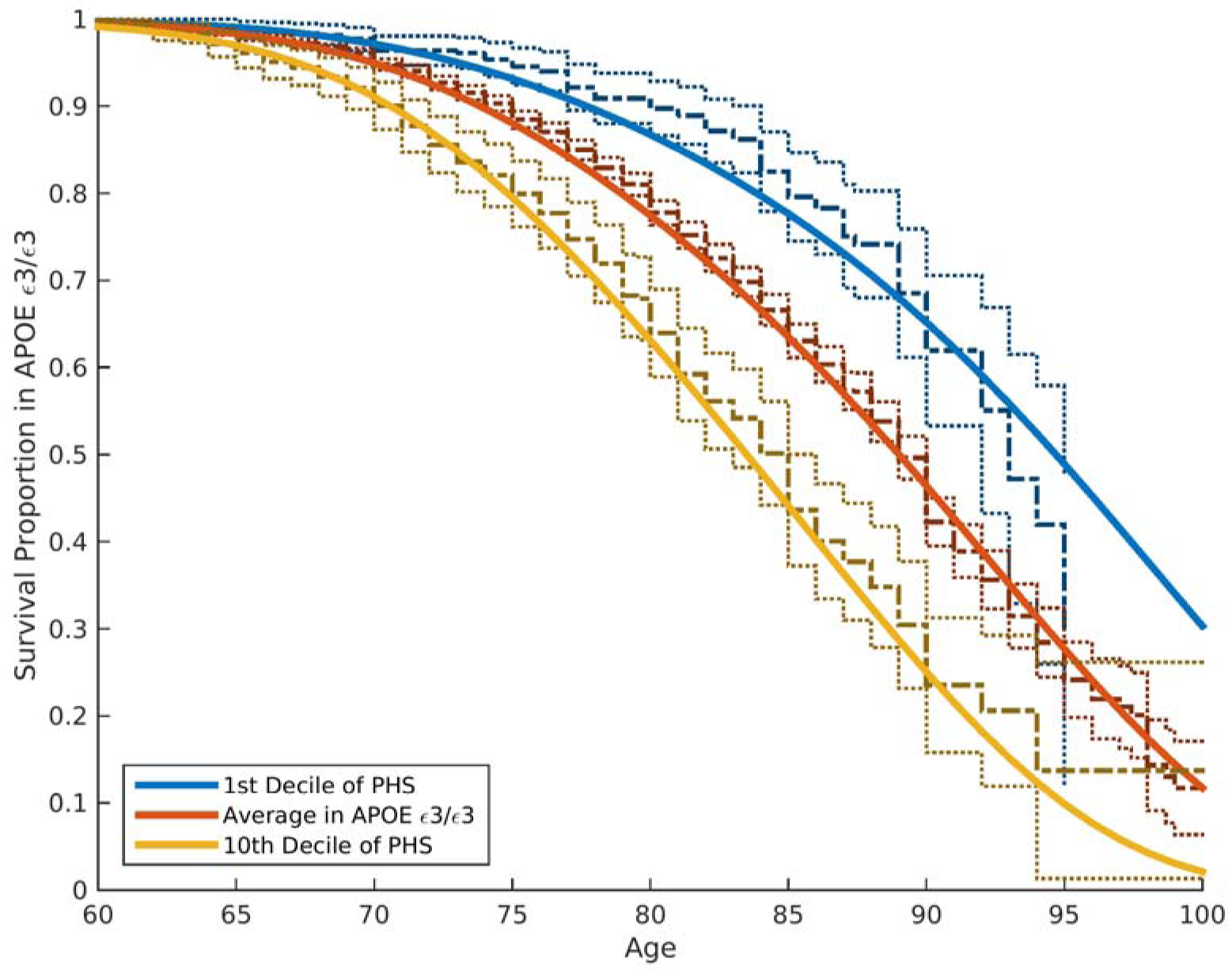
Kaplan-Meier estimates and Cox proportional model fits among *APOE* 3/3 individuals in ADGC phase 1 dataset, excluding NACC and ADNI samples.

To assess independent replication, we applied the ADGC Phase 1-trained model on independent replication samples from ADGC Phase 2. Using the empirical distributions, we found that the PHS successfully stratified individuals from independent cohorts into different risk strata (Figure 3a). Among AD cases in the ADGC Phase 2 cohort, we found that the predicted age of onset was strongly associated with the empirical (actual) age of onset (binned in percentiles, r = 0.90, p = 1.1 x 10^-26^, Figure 3b).

**Figure 3.**
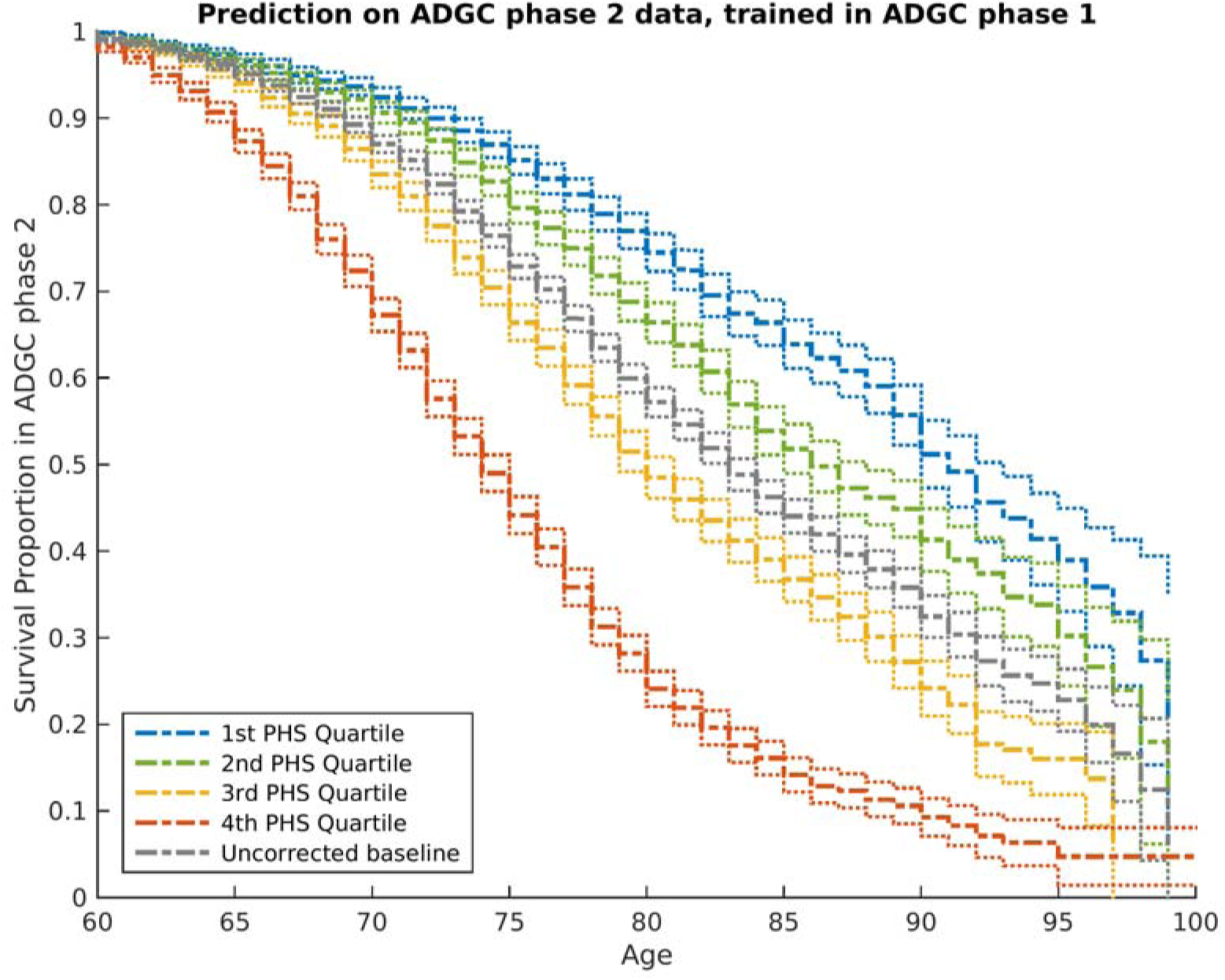
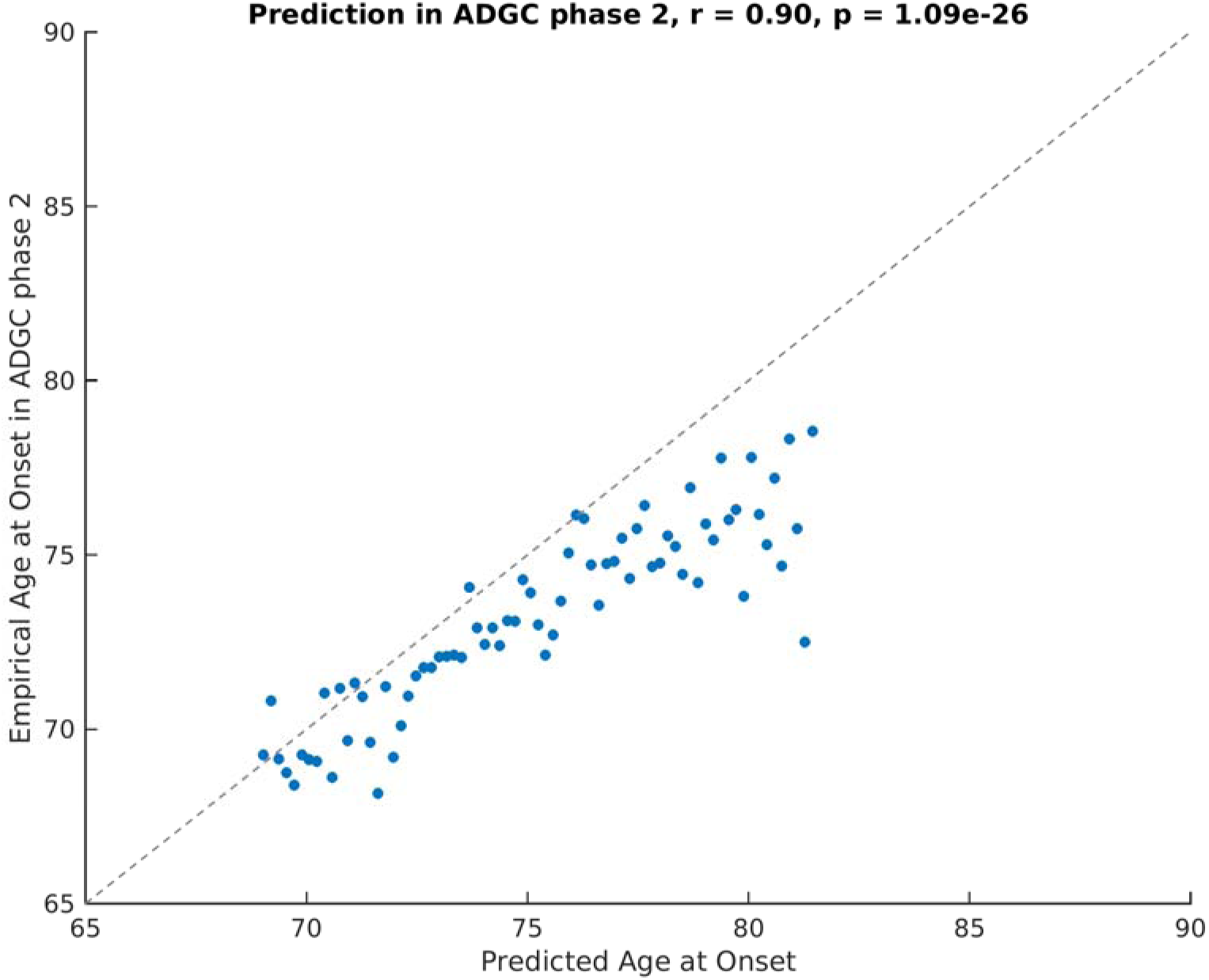
**(a)** Risk stratification in ADGC phase 2 cohort, using PHS derived from ADGC phase 1 dataset. **(b)** Predicted age of AD onset as a function of empirical age of AD onset among cases in ADGC phase 2 cohort. Prediction is based on the final survival model trained in the ADGC phase 1 dataset.

### Predicting population risk of AD onset

To evaluate risk for developing AD, combining the estimated hazard ratios from the ADGC cohort, allele frequencies for each of the AD-associated SNPs from the 1000 Genomes Project and the disease incidence in the general US population, ^17^ we generated the population baseline-corrected survival curves given an individual’s genetic profile and age (Supplemental Figures 2A and 2B). We found that the risk for developing AD as well as the distribution of age of onset is modified by PHS status (Supplemental Figures 2A,B).

Given an individual’s genetic profile and age, the corrected survival proportion can be translated directly into incidence rates (Figure 4, Table 3 and Supplemental Table 1). As previously reported in a meta-analysis summarizing four studies from the US general population, ^17^ the annualized incidence rate represents the proportion (in percent) of individuals in a given risk stratum and age, who have not yet developed AD but will develop AD in the following year; thus the annualized incidence rate represents the instantaneous risk for developing AD conditional on having survived up to that point in time. For example, for a cognitively normal 65 year-old individual in the 80^th^ percentile PHS, the incidence rate would be: 0.29 at age 65, 1.22 at age 75, 5.03 at age 85, and 20.82 at age 95 (Figure 4 and Table 3); in contrast, for a cognitively normal 65 year old in the 20^th^ percentile PHS, the incidence rate (per 100 person-years) would be 0.10 at age 65, 0.43 at age 75, 1.80 at age 85, and 7.43 at age 95 (Figure 4 and Table 3). As independent validation, we examined whether the PHS predicted incidence rate reflects the empirical progression rate (from normal control to clinical AD) (Figure 5). We found that the PHS predicted incidence was strongly associated with empirical progression rates (Cochrane Armitage trend test, p = 1.54 x 10^-10^).

**Table 3.** Predicted annualized incidence rate (per 100 person-years) by age using polygenic hazard scores.

| Age | Population Baseline* | PHS 1 percentile (95% CI) | PHS 20<sup>th</sup> percentile (95% CI) | PHS 80<sup>th</sup> percentile (95% CI) | PHS 99<sup>th</sup> percentile (95% CI) | *APOE* ε4+ (95% CI) | *APOE* ε4- (95% CI) |
| --- | --- | --- | --- | --- | --- | --- | --- |
| 60 | 0.08 | 0.02<br>(0.01,0.03) | 0.04<br>(0.01,0.08) | 0.15<br>(0.04, 0.27) | 0.61<br>(0.16, 1.06) | 0.19<br>(0.18, 0.20) | 0.06<br>(0.06, 0.7) |
| 65 | 0.17 | 0.04<br>(0.01,0.06) | 0.09<br>(0.03, 0.16) | 0.32<br>(0.09, 0.54) | 1.24<br>(0.33,2.15) | 0.38<br>(0.36, 0.40) | 0.13<br>(0.12, 0.13) |
| 70 | 0.35 | 0.07<br>(0.02,0.13) | 0.19<br>(0.05,0.32) | 0.64<br>(0.18, 1.10) | 2.53<br>(0.68, 4.38) | 0.78<br>(0.74, 0.82) | 0.26<br>(0.25, 0.27) |
| 75 | 0.71 | 0.15<br>(0.05,0.19) | 0.38<br>(0.11,0.65) | 1.30<br>(0.36,2.25) | 5.15<br>(1.38, 8.91) | 1.58<br>(1.51, 1.66) | 0.53<br>(0.52, 0.55) |
| 80 | 1.44 | 0.31<br>(0.26,0.26) | 0.77<br>(0.22,1.32) | 2.65<br>(0.74, 4.57) | 10.47<br>(2.81, 18.13) | 3.22<br>(3.06, 3.38) | 1.08<br>(1.05, 1.11) |
| 85 | 2.92 | 0.63<br>(0.19,1.07) | 1.57<br>(0.45, 2.68) | 5.39<br>(1.50, 9.29) | 21.30<br>(5.72, 36.88) | 6.55<br>(6.23, 6.87) | 2.2<br>(2.13, 2.27) |
| 90 | 5.95 | 1.28<br>(0.38,2.18) | 3.19<br>(0.91, 5.46) | 10.97<br>(3.05, 18.89) | 43.32<br>(11.63, 75.00) | 13.33<br>(12.68, 13.98) | 4.48<br>(4.34, 4.61) |
| 95 | 12.1 | 2.61<br>(0.78,4.44) | 6.48<br>(1.85, 11.10) | 22.31<br>(6.20, 38.43) | 88.11<br>(23.66, 100.00) | 27.11<br>(25.79, 28.43) | 9.1<br>(8.83, 9.38) |

\* US community-sampled population incidence proportion (% year) reported by reference 17.

**Figure 4.**
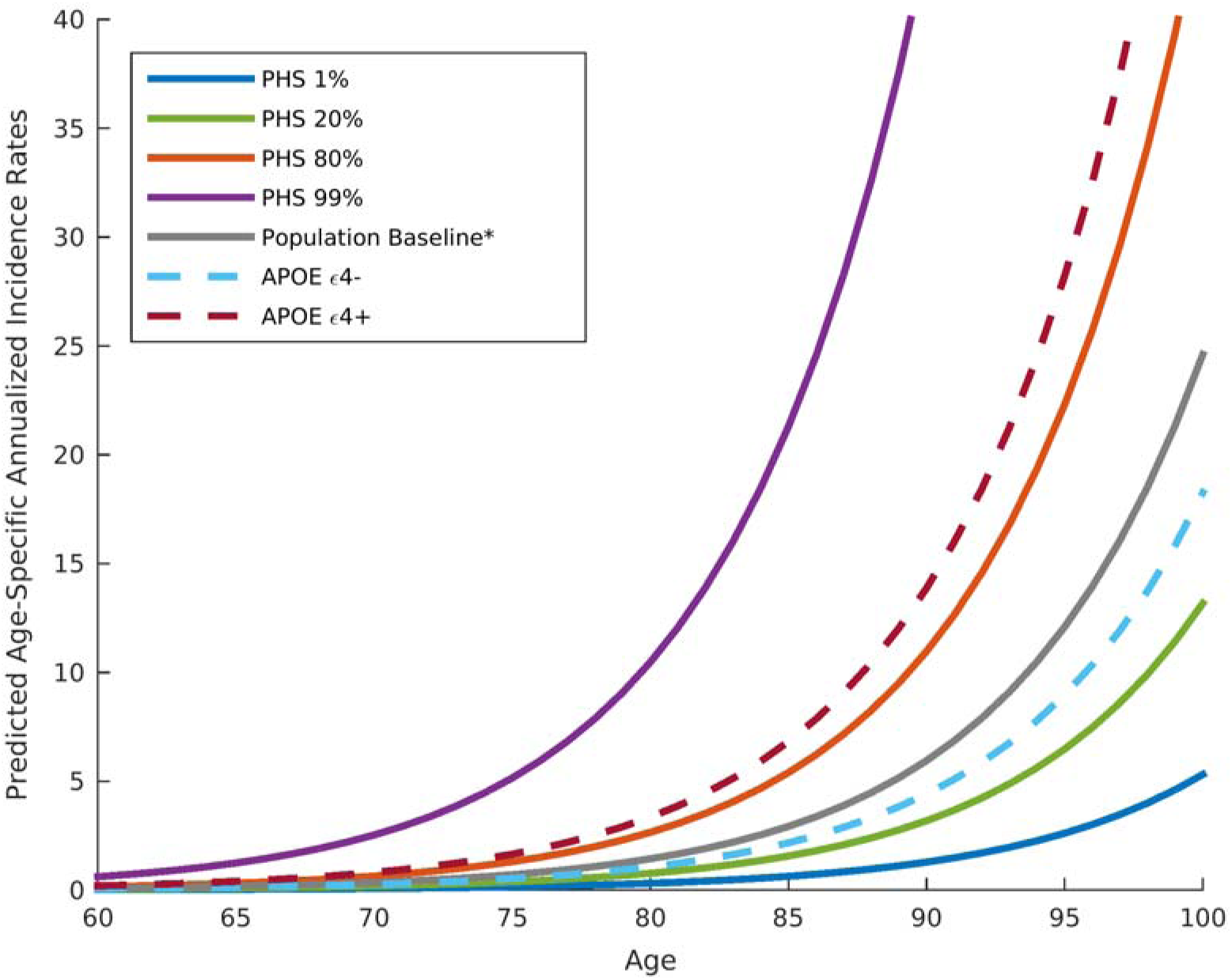
Annualized incidence rates showing the instantaneous hazard as a function of PHS percentiles and age. The gray line represents the population baseline estimate.

**Figure 5.**
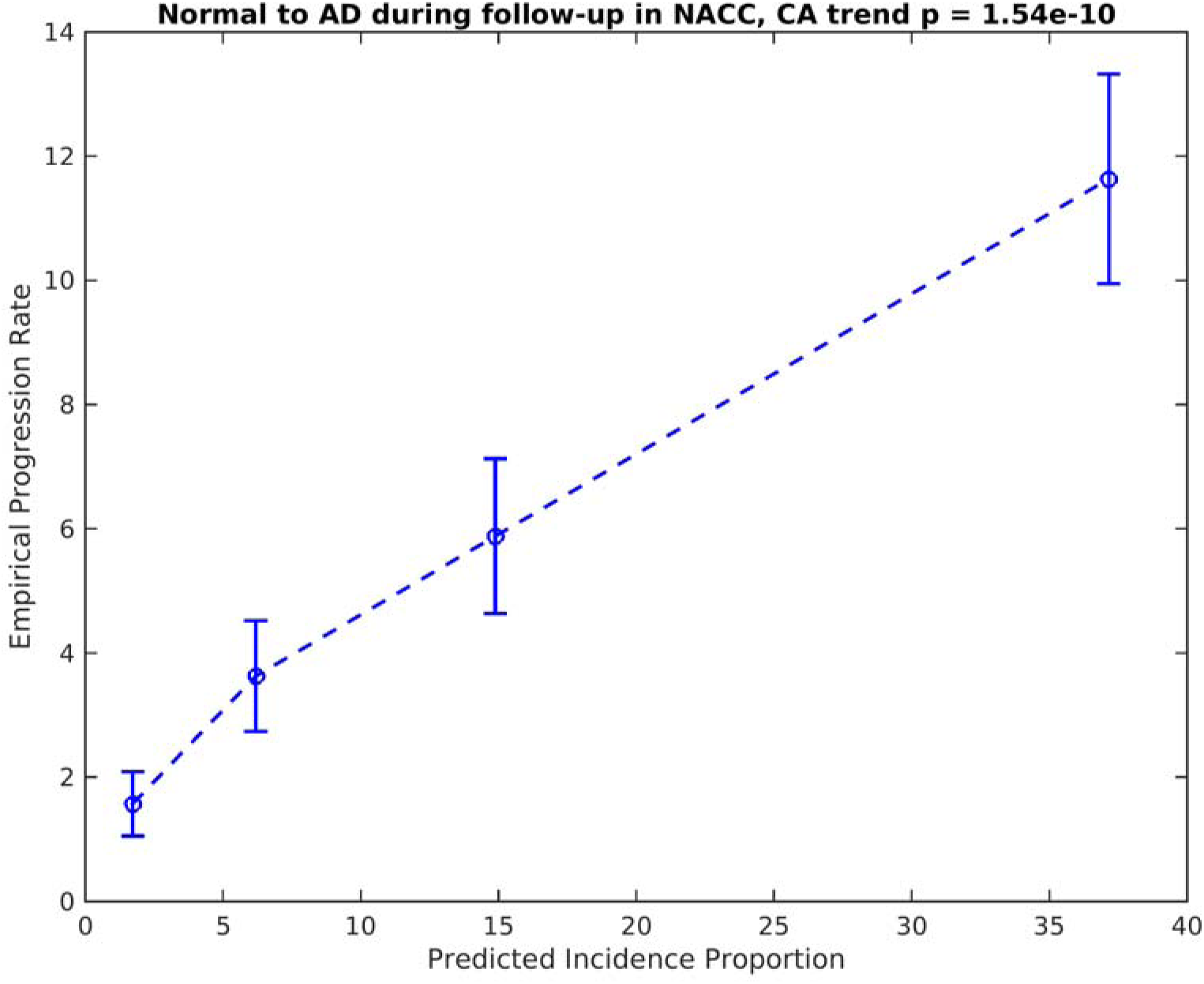
Empirical progression rates observed in the NIA ADC longitudinal cohort as a function of predicted incidence. CA = Cochrane-Armitage test

### Association with known markers of AD pathology

We found that the PHS was significantly associated with Braak stage of NFTs (β-coefficient = 0.115, standard error (SE) = 0.024, p-value = 3.9 x 10^-6^) and CERAD score for neuritic plaques (β-coefficient = 0.105, SE = 0.023, p-value = 6.8 x 10^-6^). We additionally found that the PHS was associated with worsening CDR-Sum of Box score over time (β-coefficient = 2.49, SE = 0.38, p-value = 1.1 x 10^-10^), decreased CSF Aβ_1-42_ (reflecting increased intracranial Aβ plaque load) (β-coefficient = -0.07, SE = 0.01, p-value = 1.28 x 10^-7^), increased CSF total tau (β-coefficient = 0.03, SE = 0.01, p-value = 0.05), and increased volume loss within the entorhinal cortex (β-coefficient = -0.022, SE = 0.005, p-value = 6.30 x 10^-6^) and hippocampus (β-coefficient = -0.021, SE = 0.0054, p-value = 7.86 x 10^-5^).

## DISCUSSION

In this study, by integrating AD-associated SNPs from recent GWAS and disease incidence estimates from the US population into a genetic epidemiology framework, we have developed a clinically usable, polygenic hazard score for quantifying individual differences in risk for developing AD, as a function of genotype and age. The PHS systematically modified age of AD onset, and was associated with known *in vivo* and pathologic markers of AD neurodegeneration. In independent cohorts, the PHS successfully predicted empirical (actual) age of onset and longitudinal progression from normal aging to AD. Even among individuals who do not carry the ε4 allele of *APOE* (the majority of the US population), we found that polygenic information is useful for predicting age of AD onset.

Using a case/control design, prior work has combined GWAS-associated polymorphisms and disease prediction models to predict risk for AD. ^18-19^ Rather than representing a continuous process where non-demented individuals progress to AD over time, the case/control approach implicitly assumes that normal controls do not develop dementia and treats the disease process as a dichotomous variable where the goal is maximal discrimination between diseased ‘cases’ and healthy ‘controls’. Given the striking age-dependence of AD, this approach is clinically suboptimal for predicting risk of AD. Building on prior genetic estimates from the general population, ^2, 20^ we employed a survival analysis framework to integrate AD-associated common variants with established population-based incidence ^17^ to derive a continuous measure, polygenic hazard score (PHS). From a personalized medicine perspective, for a single non-demented individual, the PHS can estimate individual differences in AD risk across a lifetime and can quantify the yearly incidence rate for developing AD.

These findings indicate that the lifetime risk of age of AD onset varies by polygenic profile. For example, the annualized incidence rates (risk for developing AD in a given year) are considerably lower for an 80-year old individual in the 20^th^ percentile PHS relative to an 80-year old in the 99^th^ percentile PHS (Figure 4 and Table 3). Across the lifespan (Supplemental Figure 2B), our results indicate that even individuals with low genetic risk (low PHS) develop AD, but at a later peak age of onset. This suggests that all individuals, irrespective of genotype, would eventually succumb to dementia if they did not die from other causes. Certain loci (including *APOE* ε2) may ‘protect’ against AD by delaying, rather than preventing, disease onset.

Our polygenic results provide important predictive information beyond *APOE*. Among *APOE* ε3/3 individuals, who constitute 70-75% of all individuals diagnosed with late-onset AD, age of onset varies by more than 10 years, depending on polygenic risk profile (Figure 2). At 60% AD risk *APOE* ε3/3 individuals in the 1^st^ decile of PHS have an expected age of onset of 85 whereas for individuals in the 10^th^ decile of PHS, the expected age of onset is greater than 95. These findings are directly relevant to the general population where *APOE* ε4 only accounts for a fraction of AD risk ^3^ and are consistent with prior work ^21^ indicating that AD is a polygenic disease where non-*APOE* genetic variants contribute significantly to disease etiology.

Using the ADGC phase 2 dataset, we found that the PHS strongly predicted actual age of AD onset in an independent sample indicating the feasibility of using PHS for diagnosing clinical AD. Within the NIA ADC sample, the PHS robustly predicted longitudinal progression from normal aging to AD illustrating the clinical value of using polygenic information to identify cognitively normal older individuals at highest risk for developing AD (preclinical AD). We found a strong relationship between PHS and increased tau associated NFTs and amyloid plaques suggesting that our genetic marker of disease risk reflects underlying Alzheimer’s pathology. The PHS also demonstrated robust associations with CSF Aβ_1-42_ levels, longitudinal MRI measures of medial temporal lobe volume loss and baseline CDR-SB score illustrating that increased genetic risk predicts clinical status and neurodegeneration *in vivo*.

From a clinical perspective, our genetic risk score, based on standard SNP chip arrays, can be used clinically for disease diagnosis, accurate identification of older individuals at greatest risk for developing AD and potentially, for informing management decisions. By providing an accurate, probabilistic assessment as to whether Alzheimer’s neurodegeneration is likely to occur, determining a ‘genomic profile’ of AD may help initiate a dialogue on future planning. Importantly, a continuous, polygenic measure of AD genetic risk may provide an enrichment strategy for prevention and therapeutic trials and could also be useful for predicting which individuals may respond to therapy. Finally, a similar genetic epidemiology framework may be useful for quantifying the risk associated with numerous other common diseases.

There are several limitations to our study. We primarily focused on Caucasian individuals of European descent. Given that AD incidence ^20^ and genetic risk ^22,23^ in African-Americans and Latinos is different than in Caucasians, additional work will be needed to develop a polygenic risk model in non-Caucasian populations. The previously reported population annualized incidence rates were not separately provided for males and females. ^17^ Therefore, we could not report PHS annualized incidence rates stratified by sex. Finally, we focused on *APOE* and GWAS-detected polymorphisms for disease prediction. Given the flexibility of our genetic epidemiology framework, it can be used to investigate whether a combination of common and rare genetic variants along with clinical, cognitive and imaging biomarkers may prove useful for refining the prediction of AD age of onset.

In conclusion, we have developed, replicated and validated a clinically useful new polygenic hazard score for quantifying the age-associated risk for developing AD. By integrating population based incidence proportion and genome-wide data into a genetic epidemiology framework, we were able to derive hazard estimates whereby an individual could calculate his/her ‘personalized’ age-specific AD risk, given genetic information. Measures of polygenic risk may prove useful for early detection, determining prognosis, and as an enrichment strategy in clinical trials.

## ACKNOWLEDGEMENTS

Drs. Rahul Desikan and Anders Dale had full access to all of the data in the study and take responsibility for the integrity of the data and the accuracy of the data analysis. Drs. Rahul Desikan (UCSF), Chun Chieh Fan (UCSD), Yunpeng Wang (UCSD and University of Oslo) and Anders Dale (UCSD) conducted and are responsible for the data analysis in this manuscript. The sources of financial and material support had no role in the design and conduct of the study; collection, management, analysis, and interpretation of the data; and preparation, review, or approval of the manuscript. We thank the Shiley-Marcos Alzheimer’s Disease Research Center at UCSD and the Memory and Aging Center at UCSF for continued support and the International Genomics of Alzheimer's Project (IGAP) for providing summary results data for these analyses. This work was supported by grants from the National Institutes of Health (NIH-AG046374, K01AG049152, R01MH100351), the Research Council of Norway (#213837, #225989, #223273, #237250/EU JPND), the South East Norway Health Authority (2013-123), Norwegian Health Association and the KG Jebsen Foundation. Please see Supplemental Acknowledgements for IGAP, NIAGADS, ADGC, ADNI and NACC funding sources.

